# “Opposite-of”-information improves similarity calculations in phenotype ontologies

**DOI:** 10.1101/108977

**Authors:** Sebastian Köohler, Peter N Robinson, Christopher J Mungall

## Abstract

One of the most important use cases of ontologies is the calculation of similarity scores between a query and items annotated with classes of an ontology. The hierarchical structure of an ontology does not necessarily reflect all relevant aspects of the domain it is modelling, and this can reduce the performance of ontology-based search algorithms. For instance, the classes of phenotype ontologies may be arranged according to anatomical criteria, but individual phenotypic features may affect anatomic entities in opposite ways. Thus, “opposite” classes may be located in close proximity in an ontology; for example enlarged liver and small liver are grouped under abnormal liver size. Using standard similarity measures, these would be scored as being similar, despite in fact being opposites.

In this paper, we use information about opposite ontology classes to extend two large phenotype ontologies, the human and the mammalian phenotype ontology. We also show that this information can be used to improve rankings based on similarity measures that incorporate this information. In particular, cosine similarity based measures show large improvements. We hypothesize this is due to the natural embedding of opposite phenotypes in vector space.

We support the idea that the expressivity of semantic web technologies should be explored more extensively in biomedical ontologies and that similarity measures should be extended to incorporate more than the pure graph structure defined by the subclass or part-of relationships of the underlying ontologies.

## Background

Ontologies have become a widely used tool to capture knowledge about objects in biology, genomics and medicine. Besides enabling knowledge integration and retrieval, they are also a widely used tool for similarity calculation between items that have been described (annotated) with classes of an ontology [1]. Ontology-based similarity measures allow non-perfect matches between ontology-classes to be quantified by incorporating the graph-structure of the ontology. Often used similarity measures included semantic similarity measures [1], cosine similarity measure, and Bayesian ontology querying [2].

Classes in an ontology are usually classified along one axis. In ontologies that try to capture phenotypic abnormalities, this axis is usually the anatomical entity in which the abnormality is seen. By this procedure, ontology classes become siblings, which would have been located in completely different parts of the ontology if the classification would be done along a different axis. An example for this are *Hyperpigmentation of the skin* (HP:0000953) and *Hypopigmentation of the skin* (HP:0001010), which are both subclasses of *Abnormality of skin pigmentation* (HP:0001000). If the classes would have been classified along the axis of qualities, the first class would belong to the *Increase of ‘something’* subontology and the latter would be in the *Decrease of ‘something’* subontology. Our hypothesis is, that such constellations lead to exaggerated similarity values for objects that do contain opposite annotations. For example, currently a gene annotated to *Hyperpigmentation* would obtain a high similarity score when compared to a gene annotated to *Hypopigmentation*.

In this paper, we explain how we added the *opposite of* information to the Phenotype and Trait Ontology (PATO) and afterwards to two large phenotype ontologies, the Mammalian Phenotype Ontology (MPO, [3]) and the Human Phenotype Ontology (HPO, [4])

Finally, we show that including this information in semantic similarity, cosine similarity, and a Bayesian algorithm can lead to an improvement in ranking objects that have been annotated with classes from the phenotype ontologies.

## Methods

The general workflow for generating *opposite_of*-relationship is depicted in Figure 1.

**Figure 1.**
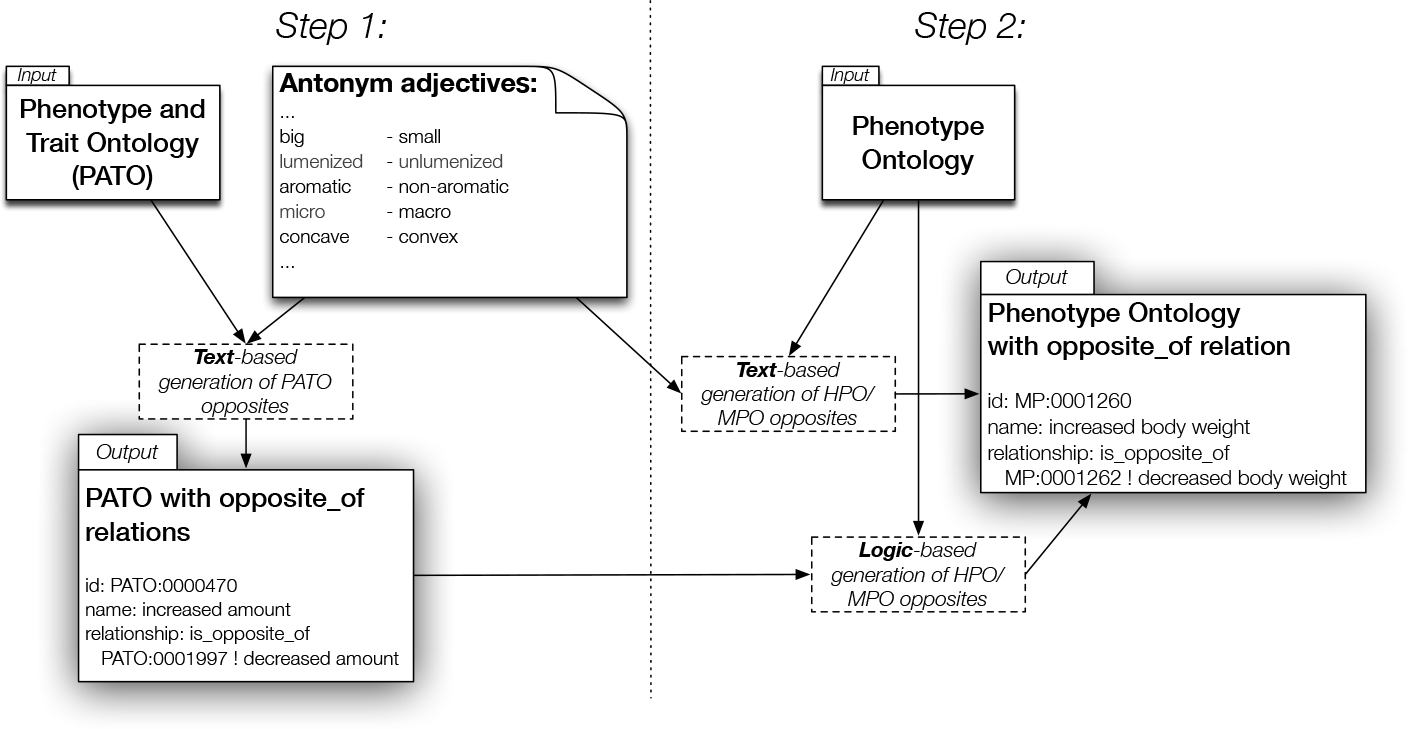
Workflow to generate opposite of-relations in phenotype ontologies. Illustration of the workflow used in this project. In Step 1, we created a list of antonym adjectives, that were used to find antonym labels in the Phenotype and Trait Ontology (PATO) and generated PATO*O* (PATO with opposites). In Step 2 we adopted two strategies to identify *opposite of*-relationships in the phenotype ontologies HPO and MPO. We applied the same strategy as in Step 1, i.e. we used the adjective list and applied string matching to find antonym phenotype labels. In parallel we used PATO*O* and the logical definitions of the phenotype classes to identify opposite classes in alogic-based procedure.

### Generation of PATO opposite of-relationships

In a first step (compare Figure 1) we compiled a list (*L*) of antonym adjective pairs that we think are good means to identify medically relevant opposite classes. Examples for antonym pairs in this list are “closed - open”, “deep - shallow”, “increased decreased”, and “huge - tiny”. We excluded pairs such as “left - right”, because we think that in a medical or biological context, very often the left part of something is not considered to be the opposite of the right part.

We performed a text-based search on all labels and synonyms in PATO, by checking if they contain an adjective from the list *L* (e.g. “increased concentration” contains “increased”). Note that we excluded “RELATED” synonyms of PATO-classes as we found those to introduce to many false positive mappings. If a label/synonym (*S*_1_) was identified, we would then check if a label/synonym (*S*_2_) in a different PATO-class for which *S*_1_ is equivalent to A(*S*_2_), where the function A(*S*) replaces the occurrence of the found adjective with its antonym counterpart from *L*. Finally, we added the mappings between opposite PATO-classes to PATO using the *opposite of*-relation. We manually fixed three cases of non-1-to-1 mappings (see https://github.com/patoontology/pato/pull/104).

### Generation of phenotype opposite of-relationships

In Step 2 (compare Figure 1), we added the *opposite of*-relation to two large phenotype ontologies. Here, we used the Human Phenotype Ontology (HPO, [4]) and the Mammalian Phenotype Ontology (MPO, [3]). We applied two different strategies to identify opposite pairs in those ontologies. Note that we performed each step separately for each phenotype ontology and did not try to identify opposite pairs between the two ontologies.

#### Phenotype opposites via text matching

We used the same procedure as for the PATO-opposite creation described before. Thus, we took the antonym list (*L*) and identified all phenotype classes in HPO/MPO using text-based search of all labels and synonyms (excluding “RELATED” synonyms).

#### Phenotype opposites via OWL-DL axioms

For each pair of phenotype classes (*P*_1_ and *P*_2_), we took the logical definitions and split it into two parts - the quality (*Q*_1_,*Q*_2_) and the remaining parts of the definition (*R*_1_,*R*_2_). Now, if *R*_1_ is equivalent to *R*_2_ and *Q*_1_ and *Q*_2_ are in a *opposite of*-relationship in PATO, we would add a *opposite of*-relationship between the phenotype classes *P*_1_ and *P*_2_.

### Evaluation of opposite of-relations in phenotype ontologies

We think that *opposite of*-relations are a useful information for several applications, e.g. clinical decision support. Here, we try to show the usefulness of the *opposite of*-relations, by showing that it improves ontology-based similarity measures. We will first define how we handle *opposite of*-relations for the subclasses of phenotype classes of an *opposite of*-mapping and then define how we integrate the *opposite of* relation into three ontology-based similarity calculation methods.

#### Inheritance of opposite of-relation to subclasses

We extended the set of *opposite of*-relation by inheriting it to the descendants of the corresponding phenotype classes. For this, we performed a depth-first-search (DFS) starting at one of the classes of the *opposite of*-mapping. The DFS stopped when a descendant class is already asserted as *opposite of* another class. We did this for both classes of the initial *opposite of*-mapping, which results in two sets. Finally, all pairs between the two sets were added as *opposite of*-pairs.

#### Including opposite of information in semantic similarity measures

We took a standard semantic similarity measure and extended it to downweight contributions to the final similarity score that were created by opposite phenotype classes. The measure is based on Resnik’s definition of information content (IC). For each class *c* in the ontology, the information content IC(*c*) is defined as the negative logarithm of the frequency of annotations to the class [5], i.e. IC(*c*) = *−* log *p_c_*, where *p_c_* is the observed frequency of items (e.g. disease or mouse genotypes) annotated to class *c* among all annotated items.

The similarity between two ontology classes is then calculated as the IC of their most informative common ancestor (MICA) [5], i.e. the common ancestor with the highest IC. For this paper, we define the semantic similarity between the annotated phenotype classes of a query (*Q*) and the annotated phenotype classes of an item (*I*) as

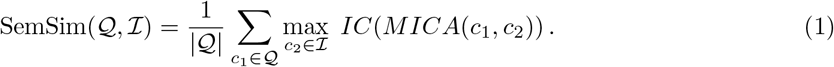

Note that *|Q|* returns the number of phenotype classes in the query *Q*.

When incorporating the information about *opposite of* relationship between two ontology classes, we introduce *Q^−^* and *I^−^* as the opposite phenotype classes of the query and of the item respectively. Note, in cases where *Q* (or *I*) do not contain any ontology class have an *opposite of*-mapping, the set *Q^−^* (or *I^−^*) may be empty.

The opposite-aware semantic similarity score (SemSim_*O*_) is then defined as:

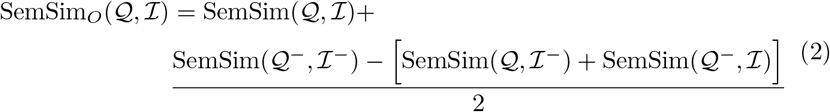

This means that we calculate SemSim as before, but we do add weight to cases where the opposites of the query and the item are similar (i.e. SemSim(*Q^−^, I^−^*)). We penalise if the query and the opposite of the item are similar (SemSim(*Q, I^−^*)) as well as if the opposite of the query and the original item are similar (SemSim(*Q^−^, I*)). After doing some evaluation tests, we chose to soften the influence of the weighting by dividing it by two.

We will later compare the ranking performance of SemSim and SemSim_*O*_.

#### Including opposite of-information in cosine similarity calculation

Another widely used similarity measure is the cosine similarity, which measures the cosine of the angle between two non-zero vectors. To apply this, the query *Q* is transferred to a vector representation ***q***, where an entry is 1 if the corresponding ontology class is present in the set and 0 otherwise. The same is done for the item *I*, i.e. it is transferred to ***i***. Note that during the transformation, all the ancestors of the classes in *Q* and *I* are set to 1 as well. The cosine similarity is then defined as

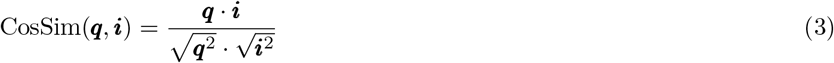

In order to include *opposite of*-information into the cosine similarity, we obtain the opposite query vector (***q**^−^*) and opposite item vector (***i**^−^*). The opposite-aware cosine similarity is then defined as:

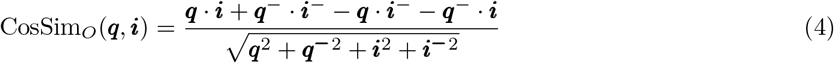

#### Including opposite_of-information in Bayesian ontology query algorithms

In order to test the effect of *opposite of*-information on Bayesian ontology querying, we adapted Algorithm 1 from the paper of Bauer et al. [2]. For a given set *S*, *S_anc_* denotes the set of all classes in *S* and all the induced ancestors of these ontology classes. The set *O* denotes the set of all terms in the corresponding ontology.

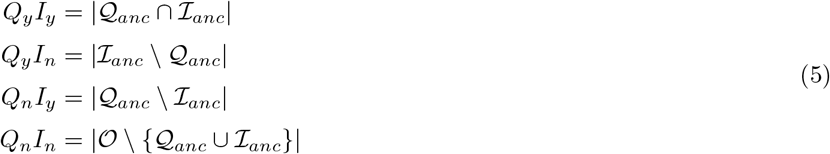

In the extended opposite-aware algorithm we are going to replace two of these numbers(*Q_y_ I_n_* and *Q_n_I_y_*). Again, we define *Q^−^_anc_* as the opposite phenotype classes of *Q_anc_*.

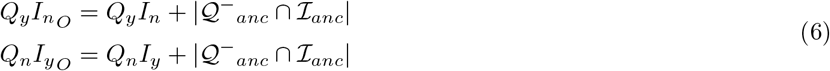

This means that we determine the number of *opposite of* mappings between the query and the item and use this number to increase the values *Q_y_ I_n_* and *Q_n_I_y_*.

As in Bauer et al. [2] we use a set of false positive rates (FPR) and a set of false negative rates (FNR), which we set to FPR = {1 *∗ e^−^*^10^, 0.0005, 0.001, 0.005, 0.01*}* and FNR = {1 *∗ e^−^*^10^, 0.005, 0.01, 0.05, 0.1, 0.2, 0.4, 0.8, 0.9*}*. The score for each item is then determined as

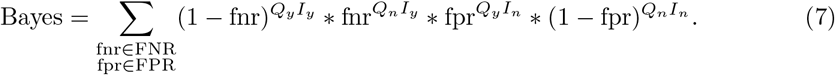

For the opposite-aware version of this score we apply the equation

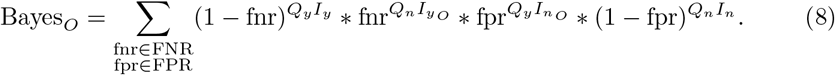

We will later compare the ranking performance of Bayes and Bayes_*O*_.

#### Test by ranking items using phenotype ontologies

For the tests in the HPO we used an HPO OBO version from January 2017 together with annotations of diseases (items) from OMIM [6], Orphanet [7], and DECIPHER [8]. For the test in the MPO we downloaded the ontology (data-version: releases/2017-01-06) as well as the annotations of genotypes (items, file: MGI PhenoGenoMP.rpt) on January 13th of 2017.

Each item (human disease or mouse model) that was annotated with at least three classes of the phenotype ontologies (i.e. *|I| ≥* 3) was considered for our simulations. At first we generate a query consisting of phenotype ontology classes that are related to, but not exactly the same as, the originally annotated classes. For this we took all phenotype classes in *I*, mapped annotated classes to one of their ancestors (1), added randomly selected phenotype classes (2), and finally selected a subset of these classes as the query.

For (1), we chose to replace 60 % of the terms to be replaced by one of its ancestor classes. We did not allow terms to be mapped to a very general class, i.e. “Phenotypic abnormality” (HP:0000118) / “mammalian phenotype” (MP:0000001) and all of its direct subclasses like “Abnormality of the musculature” (HP:0003011) / “muscle phenotype” (MP:0005369). For (2) we determined the size of query set and added 30 % randomly selected classes. In (3) we set the number *q* of phenotype classes to be selected between 5 and 9 (both inclusive). We then randomly selected *q* phenotype classes out of the set generated in (1) and (2) as the query.

Afterwards we ranked all annotated items (i.e. all diseases or mouse genotypes) by similarity to the generated query using the six mentioned methods; SemSim, SemSim_*O*_, CosSim, CosSim_*O*_, Bayes, and Bayes_*O*_. We record the rank of the initial item in the ranked list. By doing this for all items included in the simulation, we obtain a list of ranks on which we can compute several metrics (see Results).

## Results

### Generation of opposite of-relations

In Step 1 (see Figure 1) we created a list of 143 antonym adjectives. The full list is available on our GitHub repository at https://github.com/phenomics/phenopposites. We used this list to search for opposite ontology classes in the Phenotype and Trait Ontology (PATO). Using a string-matching strategy, we generated a set of 211 opposite relations in PATO. We removed three cases where non 1-to-1 mappings were generated (see https://github.com/patoontology/pato/pull/104), such that we finally have added 208 mappings, i.e. 416 “relationship: is opposite of” lines.

In Step 2 (see Figure 1) we used two different strategies to add *opposite of* relations between phenotype classes in the Mammalian and Human Phenotype Ontology (MPO, HPO). In one strategy, we took the same list of antonym adjectives as before to perform a text-based search with the primary labels and synonyms of the phenotype classes. In another strategy, we used the PATO ontology and the logical definitions of the phenotype classes. The logical definitions are OWL-DL axioms that basically define a phenotype (e.g. “short tibia”) using the intersection of a quality (from PATO, e.g. “short”) and a bearer, where the quality inheres in (e.g. “tibia bone”). We would define two phenotype classes as opposites, if the bearer is equivalent, but the their qualities are *opposite of* each other in PATO (see Methods).

The results of identified opposite phenotype classes for HPO and MPO are shown in Table 1. For MPO, we could identify approximately three times more *opposite of*-relationships (1517) than for HPO (436), which is probably caused by the fact that MPO has a much more regular structure, especially in their naming conventions, whereas HPO uses more often medical terminology which leads to much more heterogeneity in the labels and synonyms. The list of *opposite of*-relationships between classes of HPO and MPO is available on our GitHub repository (https://github.com/phenomics/phenopposites) and we plan to add those relations to the core ontologies in one of the next releases.

**Table 1.** Number of *opposite of*-mappings created in phenotype ontologies based on text-based search (*T*) and logical definition based search (*L*). The table also shows how many mappings were found only with one method (*T\L* and *L\T*), by both methods (*T ∩L*), and by at least one method (*T ∪L*).

| Ontology | $\vert \mathcal{T}\vert $ | $\vert \mathcal{L}\vert $ | $\vert \mathcal{T} \setminus \mathcal{L}\vert $ | $\vert \mathcal{L} \setminus \mathcal{T}\vert $ | $\vert \mathcal{T} \cap \mathcal{L}\vert $ | $\vert \mathcal{T} \cup \mathcal{L}\vert $ |
| --- | --- | --- | --- | --- | --- | --- |
| HPO | 236 | 384 | 200 | 52 | 184 | 436 |
| MPO | 968 | 1476 | 549 | 41 | 927 | 1517 |

For the text-matching, we added a few exceptions that we did not consider as phenotype opposites. For example, we found that abnormalities of the “large vessels” or the “large intestine” are not the opposite of abnormalities of “small vessels” or the “small intestine”. Similarly, we excluded abnormalities in “CD11b-high” because the text-based search would identify abnormalities in “CD11b-low” cells as opposite.

### Performance of opposite-aware similarity measures

We took each item that has been annotated with at least three classes from HPO or MPO as a test case. This corresponds to 9424 human diseases and 11,876 mouse genotypes. We generated a query from the given annotations as described in the Methods section. We used the query to calculated the similarity of the query to all items in the corresponding dataset and recorded the rank of item that was used to generate the query in the beginning. We did this for three similarity measures that are described in the methods section and compared them to the opposite-aware counterpart. (SemSim vs. SemSim_*O*_, CosSim vs. CosSim_*O*_, and Bayes vs. Bayes_*O*_). At first we looked if including the *opposite of*-information does have an effect on the ranking performance, and if an effect is positive (smaller rank) or negative (higher rank). In Figure 2 one can see that including *opposite of*-information does have more often an effect for MPO (average 61 %) than for HPO (average 51 %), which is likely caused by the fact we have substantially more *opposite of*-relations in MPO (see Table 1). One can also see that in cases where the *opposite of*-information has an effect, it is very often has a positive effect. Only for SemSim_*O*_ the effect is negative in 11 % (HPO) and in 18 % (MPO) of the cases. In Figure 3 we show the distribution of changed ranks for the positive and negative effects. One can clearly see that the cosine similarity measure does profit the most from the inclusion of *opposite of* information. For the Bayesian method a strong positive influence can be seen, but most often the rank improves by about one to five positions.

**Figure 2.**
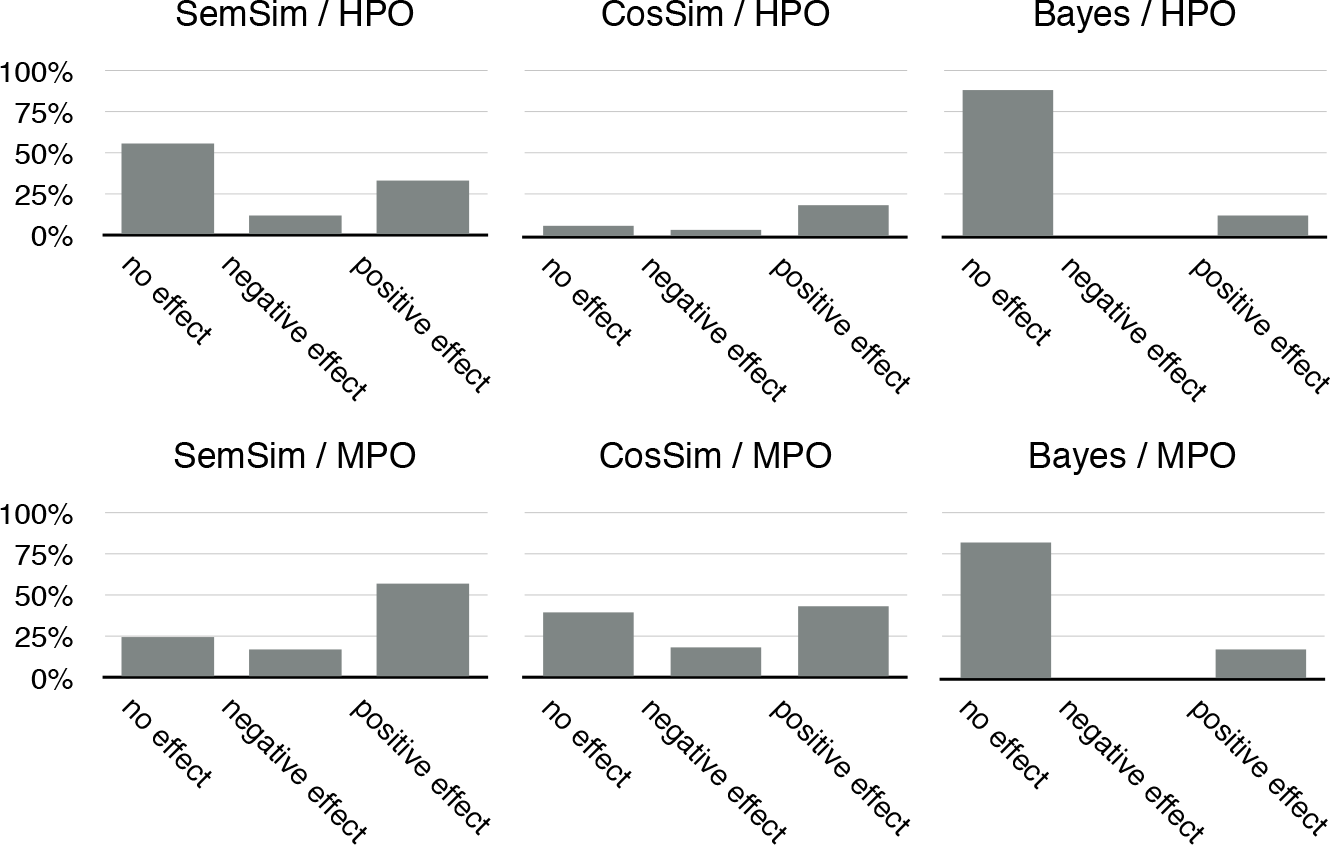
**Figure 2 Distribution of effect vs no effect on ranking items annotated with HPO or MPO.** The barplot shows how often the introduction of *opposite of*-information in the similarity measures (semantic similarity (SemSim), cosine similarity (CosSim), and Bayes) has an effect, and if this effect is positive or negative (i.e. leads to better or worse rank of the sought item). As expected, in HPO we have less often changes in the ranking, as there are currently fewer *opposite of*-relationships discovered in HPO. With the exception of SemSim, the *opposite of*-information has almost always a positive effect on ontology-based rankings. In SemSim the effect is negative in 11 % (HPO) and in 18 % (MPO) of the cases.

**Figure 3.**
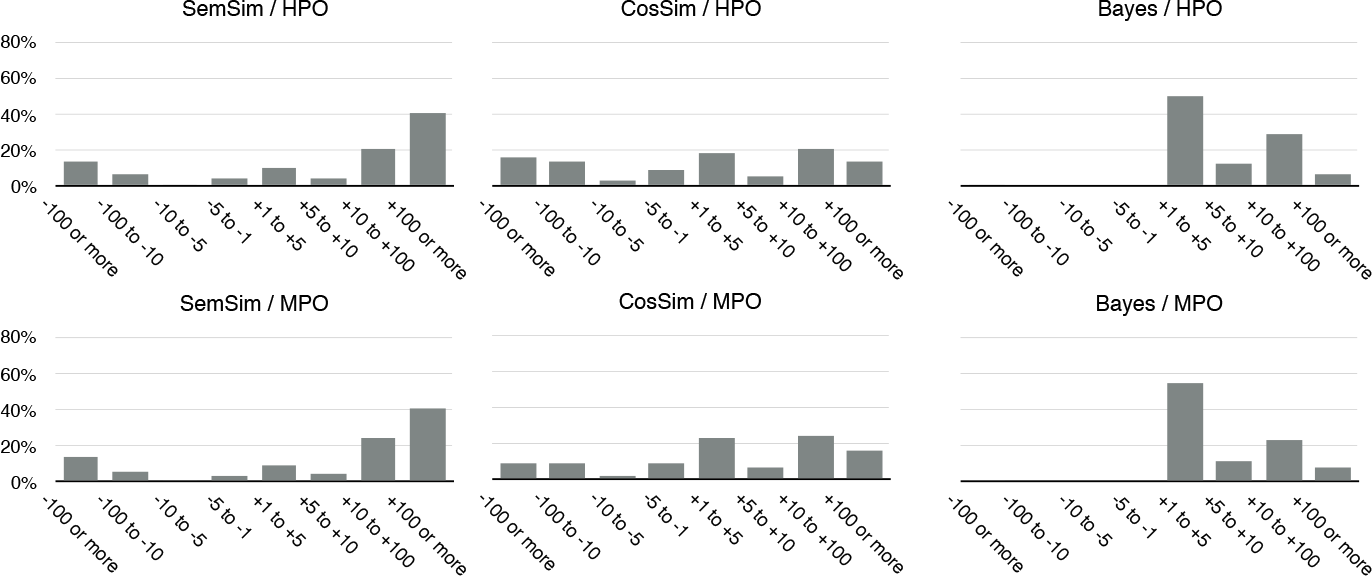
**Rank-changes when opposite of-information does change the rank of sought items.** The histograms show the distribution of the effects on rankings in cases where the inclusion of *opposite of*-information has an effect on the rank of the sought after item. The histograms are created for all combinations of ontologies (HPO, MPO) and algorithms (semantic similarity (SemSim), cosine similarity (CosSim) and Bayes) tested in this project.

Precision/recall (PR) curves are visual representations of the performance of a model in terms of the precision and recall statistics. For different thresholds it plots the actual precision (y-axis) and recall (x-axis) points and connects them by a line. An important measure is the area under this curve, which is to be maximised. In Table 2 we list the values of the area under the precision-recall curve for the different methods. One can again see that including the *opposite of*-relation improves these values as well, for semantic similarity, cosine similarity and Bayesian ontology querying. For Bayes the increase is rather small (1.01 for HPO, 1.02 for MPO). for SemSim it increases by a factor of 1.4 (HPO) and 2.9 (MPO). For CosSim the area under PR curve increases by a factor of 1.23 (HPO) and 1.3 (MPO). We do however note that it is much harder for Bayes to improve already high values of 0.23 (HPO) or 0.34 (MPO) - a performance that none of the other methods achieves.

**Table 2.** Performance measured by the area under the precision-recall (PR) curve for three tested algorithms - with and without inclusion of *opposite of*-information.

| Ontology | Algorithm | Area under PR curve |
| --- | --- | --- |
| HPO | SemSim | 0.031 |
|  | SemSim<sub>O</sub> | 0.042 |
|  | CosSim | 0.152 |
|  | CosSim<sub>O</sub> | 0.187 |
|  | Bayes | 0.228 |
|  | Bayes<sub>O</sub> | 0.23 |
| MPO | SemSim | 0.013 |
|  | SemSim<sub>O</sub> | 0.038 |
|  | CosSim | 0.213 |
|  | CosSim<sub>O</sub> | 0.275 |
|  | Bayes | 0.342 |
|  | Bayes<sub>O</sub> | 0.348 |

## Discussion

We have introduced *opposite of*-relationships between classes of three major phenotyping ontologies, PATO, MPO, and HPO. We used a text-based strategy, for which we created a list of antonym adjectives. We also used a strategy in which we inferred *opposite of* relationship based on the logical definition of phenotype classes in combination with the *opposite of*-relations we created in PATO before. We have added 208 *opposite of*-relations to PATO and 436 and 1517 *opposite of*-relations to HPO and MPO respectively. We are planning to maintain these automatically generated relationships by both manual curation in combination with further execution of the software to automatically (i.e. Step 2 in Figure 1) to suggest additions.

We tested if the performance of ranking items based ontological similarity measures is influenced by the inclusion of *opposite of* information. For this we modified a standard semantic similarity measure, a cosine similarity measure and a Bayesian approach to incorporate this information.

We do not claim that our approaches to integrate *opposite of* information in similarity calculation methods (SemSim_*O*_, CosSim_*O*_, Bayes_*O*_) are the best possible way. We rather wanted to show, that *opposite of*-information is relatively easy to generate automatically (especially, but not only for phenotype ontologies) and that incorporation of this information in such algorithms is possible and useful. We aim to generate more sophisticated statistical models to incorporate more semantic relationships in the future.

Related work was done by Ferreira et al. [9]. They investigated the usage of existing *disjoint with*-axioms for improved calculation of the MICA between two ontology classes. First, we note that we purposely decided to use the *opposite of* relation instead of *disjoint with*-axioms. In our use-case, using disjointness makes only sense when the phenotype ontology would exclusively be used to describe single individuals. For example, a single individual can never be smaller and bigger at the same time, and *disjoint with*-axioms can help finding such inconsistent annotations. However, in reality, phenotype ontologies are very often used to describe groups of individuals, such as the set of all patients with a particular disease. Thus, it is indeed correct that the HPO team has annotated “Cerebral creatine deficiency syndrome-1” (OMIM entry 300352) with both “Tall stature” (HP:0000098) and “Short stature” (HP:0004322), because it was found that two patients were tall and thin, and three had short stature [10]. These two classes (HP:0000098 and HP:0004322) are now in a *opposite of*-relationship in HPO. If we would have added the information about opposite phenotype classes as *disjoint with*-axioms, the knowledge base provided by HPO-consortium would become inconsistent. Also, Ferreira et al. [9] did not show how the inclusion of *disjoint with*-axioms affects semantic similarity between two sets of ontology classes, but rather concentrated on a sophisticated procedure to include disjointness in the calculation of the IC of the MICA (compare Equation 1).

As mentioned before, the development of more sophisticated algorithms is subject to future research. We think that considering the full logical formalism of biomedical ontologies in similarity measure has the potential to further strengthen the role of ontologies in different areas of biomedical research [11, 9].

## Abbreviations

HPO: Human Phenotype Ontology, MPO: Mammalian Phenotype Ontology, PATO: Phenotype and Trait Ontology, SemSim: semantic similarity, CosSim: Cosine similarity, DFS: depth first search, PR: precision-recall

## Author’s contributions

SK, PNR, and CM designed the study. SK performed experiments, and analysed the data. SK, PNR, and CM wrote the paper.

## Acknowledgements

This work has been funded by E-RARE project Hipbi-RD (Harmonising phenomics information for a better interoperability in the RD field, http://www.hipbi-rd.net, 01GM1608)

